# CyTOF-based profiling of circulating tumor cells predicts aggressiveness and therapy response in SCLC liquid biopsies at a personalized level

**DOI:** 10.64898/2025.12.16.694669

**Authors:** Mukulika Bose, Cole Ruoff, Shafqat Ehsan, C. Allison Stewart, Alberto Duarte, Sayantan Bhattacharyya, Jiaqi Li, Benedict Anchang, Thomas Welte, Ashley Victorian, Fabiola E. Lujan, Lixia Diao, Jing Wang, Ken Chen, Bingnan Zhang, Runsheng Wang, Luisa Solis Soto, Alejandra G. Serrano, Robert Cardnell, Carl M. Gay, Lauren A. Byers, Loukia G. Karacosta

## Abstract

Small-cell lung cancer (SCLC) is an aggressive neuroendocrine carcinoma characterized by high numbers of circulating tumor cells (CTCs). We applied CyTOF and a 20-marker antibody panel to detect and phenotype CTCs directly in liquid biopsies of 51 SCLC patients (treatment-naïve, chemotherapy and immunotherapy-treated, and tarlatamab-treated), of which a subset were longitudinally tracked. Unsupervised clustering revealed distinct cell populations enriched in patient liquid biopsies compared to those from healthy donors. Further analysis identified CTC populations of the three established SCLC subtypes driven by the high expression of ASCL1, NeuroD1, and POU2F3 transcription factors respectively. Significant differences in CTC EMT markers, established therapeutic targets (e.g. DLL3), and subtype heterogeneity were observed between naïve versus treated samples. Changes in subtype proportions were observed in longitudinally tracked samples in both treatment modalities. Our study demonstrates the utility of CyTOF for high-resolution CTC profiling, offering dynamic insights into CTC heterogeneity, treatment response, and resistance mechanisms.

**Highlights:**

- CTCs can be detected, subtyped and phenotyped in SCLC liquid biopsies using CyTOF
- CTC subtypes and EMT states are differentially associated with treatment modality
- CTC DLL3 levels and epithelial features increase following anti-DLL3 BiTE therapy
- CyTOF CTC subtyping can predict disease aggressiveness
- Longitudinal tracking reveals CTC plasticity and therapy response correlations

**Graphical Abstract:** 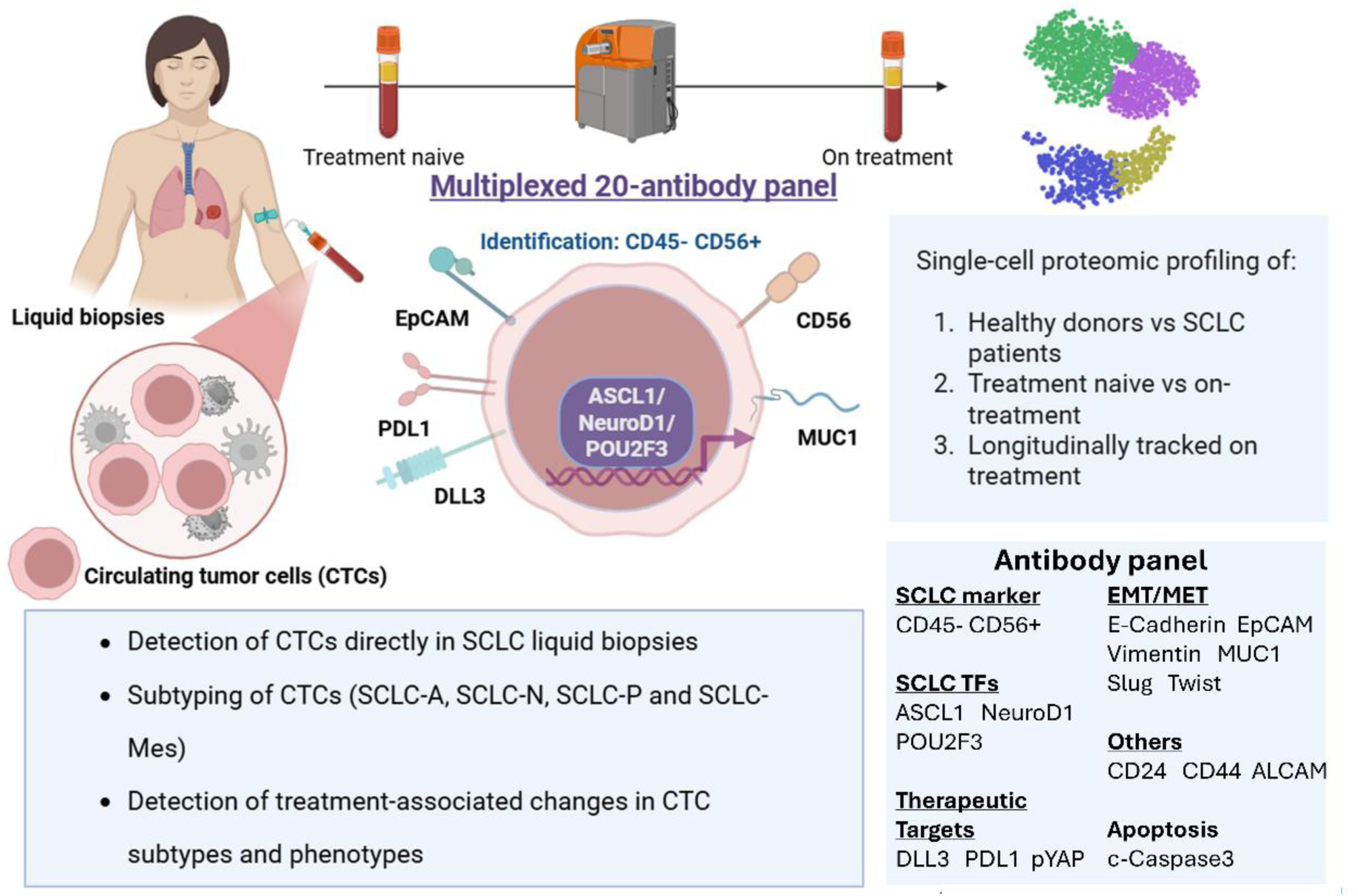

## 1. Introduction

Small cell lung cancer (SCLC) is an aggressive neuroendocrine carcinoma characterized by rapid growth, early dissemination, and poor prognosis. It forms about 15% of all lung cancer cases with a dismal 5-year survival rate of 9% [1]. Despite initial sensitivity to chemotherapy and radiotherapy, most patients inevitably experience relapse within months, highlighting the urgent need for improved prognostic tools and therapeutic strategies. SCLC typically presents as centrally located masses in the lungs, making surgical resection uncommon and tissue biopsies limited in both quantity and quality. Biopsies are also often limited in SCLC due to the urgency of treatment. The American College of Chest Physicians (ACCP) recommends relying on less invasive methods (e.g., bronchoscopy, cytology) for diagnosis when biopsy of a solitary extra-thoracic lesion is not feasible. Similarly, the National Comprehensive Cancer Network (NCCN) advises that treatment for SCLC should not be delayed more than one week for staging or procedures, due to the aggressive nature of the disease [2].This scarcity of solid tumor specimens poses significant challenges to molecular and cellular characterization [3]. Recently, liquid biopsy has emerged as an alternative, minimally invasive approach for tumor profiling, offering several advantages over traditional tissue biopsies. By analyzing components shed into the bloodstream - such as circulating tumor DNA (ctDNA), exosomes, and circulating tumor cells (CTCs), liquid biopsies allow for real-time monitoring of tumor cell dynamics, clonal evolution, and treatment response [4–7]. Among these analytes, CTCs represent a particularly valuable resource, especially in SCLC where their frequency in peripheral blood is relatively high compared to other solid tumors [8]. CTCs provide a window into the viable tumor cell population in circulation, and their phenotypic heterogeneity may reflect mechanisms of metastasis, therapeutic resistance, and disease progression [9–11].

While high CTC counts are linked to poor prognosis in SCLC, they do not consistently predict treatment response, highlighting the need for deep phenotyping beyond enumeration [12]. To fully harness the potential of CTCs as dynamic biomarkers, high-dimensional single-cell technologies are essential [13]. Building on our previous work applying CyTOF for profiling non-small cell lung cancer (NSCLC) cell lines, tumors, and pleural effusions [14] [15], we extended this approach to CTCs in SCLC. Our CyTOF approach is uniquely suited to capture the full phenotypic complexity of CTCs, including subtype-defining transcription factors, epithelial-mesenchymal transition (EMT) states, therapeutic targets, and protein markers associated with drug resistance (specifically platinum-based chemotherapy, immunotherapy and the recently FDA-approved anti-DLL3 bi-specific T-cell engager or BiTE, called tarlatamab). By applying this technology to SCLC liquid biopsies, we aimed to uncover clinically meaningful subpopulations that may inform prognosis and guide real-time treatment decisions - ultimately contributing to the development of more personalized and adaptive therapeutic strategies for this aggressive malignancy with limited targeted therapy options.

## 2. Results

### 2.1 CyTOF-based detection and characterization of CTCs in SCLC liquid biopsies

To detect and interrogate CTC phenotypes in SCLC liquid biopsies, we optimized a CyTOF workflow for whole blood processing for isolating peripheral blood mononuclear cells (PBMCs), antibody staining, and high-dimensional analysis. Peripheral blood samples from 51 SCLC patients and 19 healthy donors were processed enabling direct comparison of circulating cell populations to establish baseline SCLC-specific marker expression and cell phenotypes [**Table 1 (Abacus plot)**]. We designed a custom 20-marker antibody panel targeting SCLC-related biomarkers and therapeutic targets (e.g. DLL3), the three SCLC subtype-defining transcription factors NeuroD1, ASCL1, and POU2F3[16] and EMT-related markers[14] (**Supplementary Table 1**). To identify CTC-like populations, we first used a CD45-CD56+ gating scheme to exclude immune cell populations **(Supplementary Figure 1A)**. CD56 is expressed in ∼90% of SCLC cells [17]-a feature we confirmed in SCLC cell lines, whereas EpCAM was expressed only on a subset (∼30%) of SCLC cells, further validating the use of CD56 in our gating scheme, as opposed to an epithelial marker such as EpCAM which is commonly used for epithelial CTC isolation methodologies [18] (**Supplementary Figure 1B**). To optimize and validate SCLC subtype-defining transcription factor expression, we utilized reference SCLC cell lines of established biomarker expression as positive and negative controls (**Supplementary Figure 1C**). Further details on sample processing, staining optimization, technical replicability, and batch effect correction can be found in the Methods section and **Supplementary Figures 1D-F.**

**Table 1:**
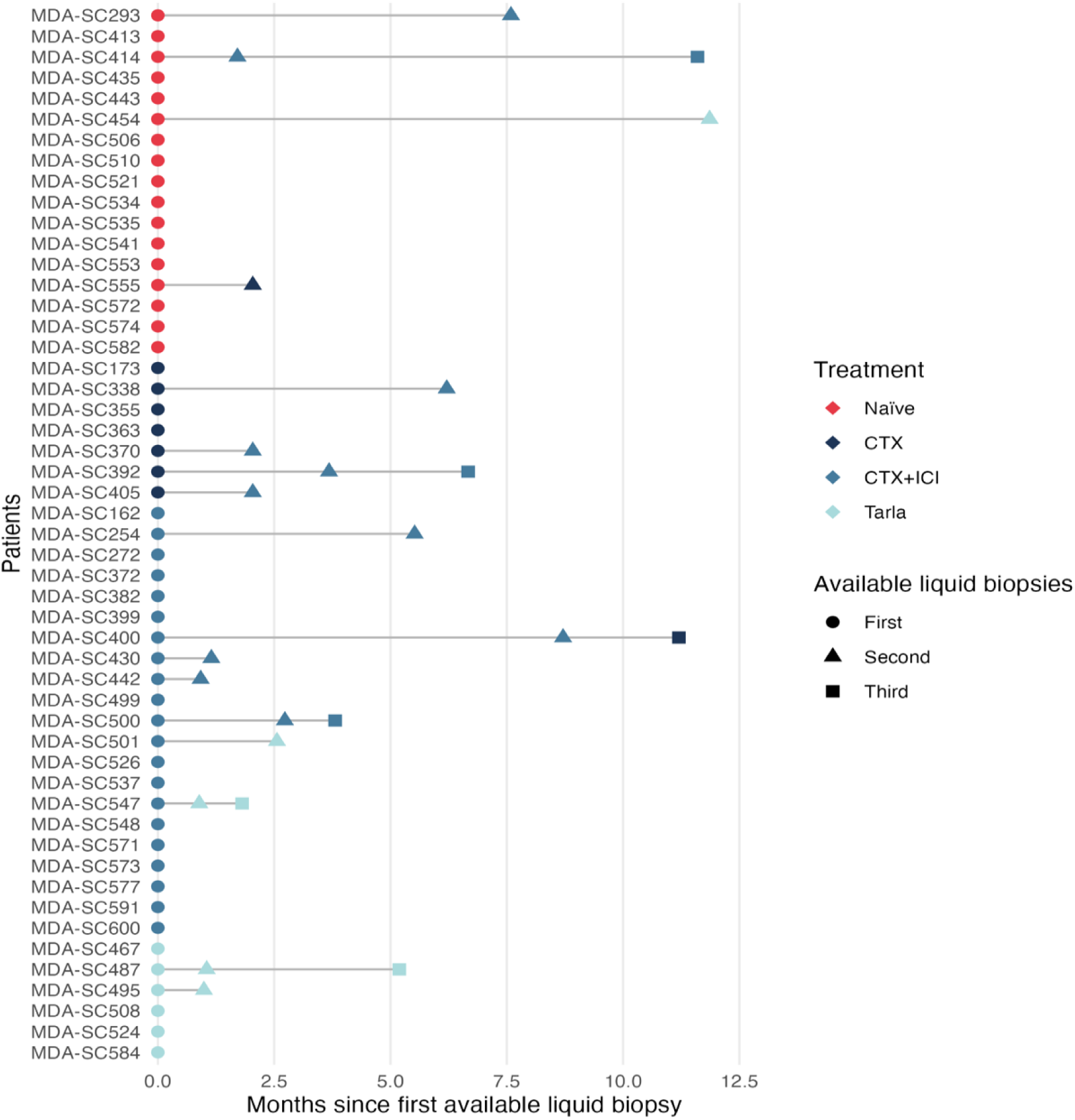
Abacus plot showing patient and liquid biopsy samples collected for CyTOF Analysis Timeline of liquid biopsy collection for patients stratified by treatment type. Each patient is represented on the y-axis, with time (months) since the first available liquid biopsy on the x-axis. Symbols indicate the sequence of liquid biopsies collected: circles for the first, triangles for the second, and squares for the third blood draw. Colors denote treatment groups: red for treatment-naïve, dark blue for chemotherapy (CTX), light blue for chemotherapy plus immune checkpoint inhibition (CTX+ICI), and turquoise for patients treated with tarlatamab (Tarla). This timeline illustrates the sampling schedule across different treatment regimens and patient follow-up durations.

To filter out non-CTC like cell populations *in silico*, we implemented a two-step (primary and secondary) clustering approach (**Supplementary Figure 2A**). First, we performed unsupervised clustering on pooled CD45-CD56+ cells from 19 blood samples from healthy donors and 75 liquid biopsies from 51 SCLC patients (**Table 1**) using the FlowSOM algorithm (**Figure 1A**). CD45-CD56+ cells from both healthy donors and patients grouped into eight primary clusters (p1-p8) (**Figure 1A** and **Supplementary Figure 2B**). Primary clusters p6, p7, and p8 were more abundant in SCLC patient samples compared to healthy donors, with p6 and p7 showing statistically significant enrichment. These three primary clusters were collectively designated as cancer-enriched clusters (**Figure 1B**). Cluster p8 was included as a cancer-enriched cluster despite the lack of statistically significant enrichment, given that it almost entirely originated from the liquid biopsy of a single SCLC patient.

**Figure 1.**
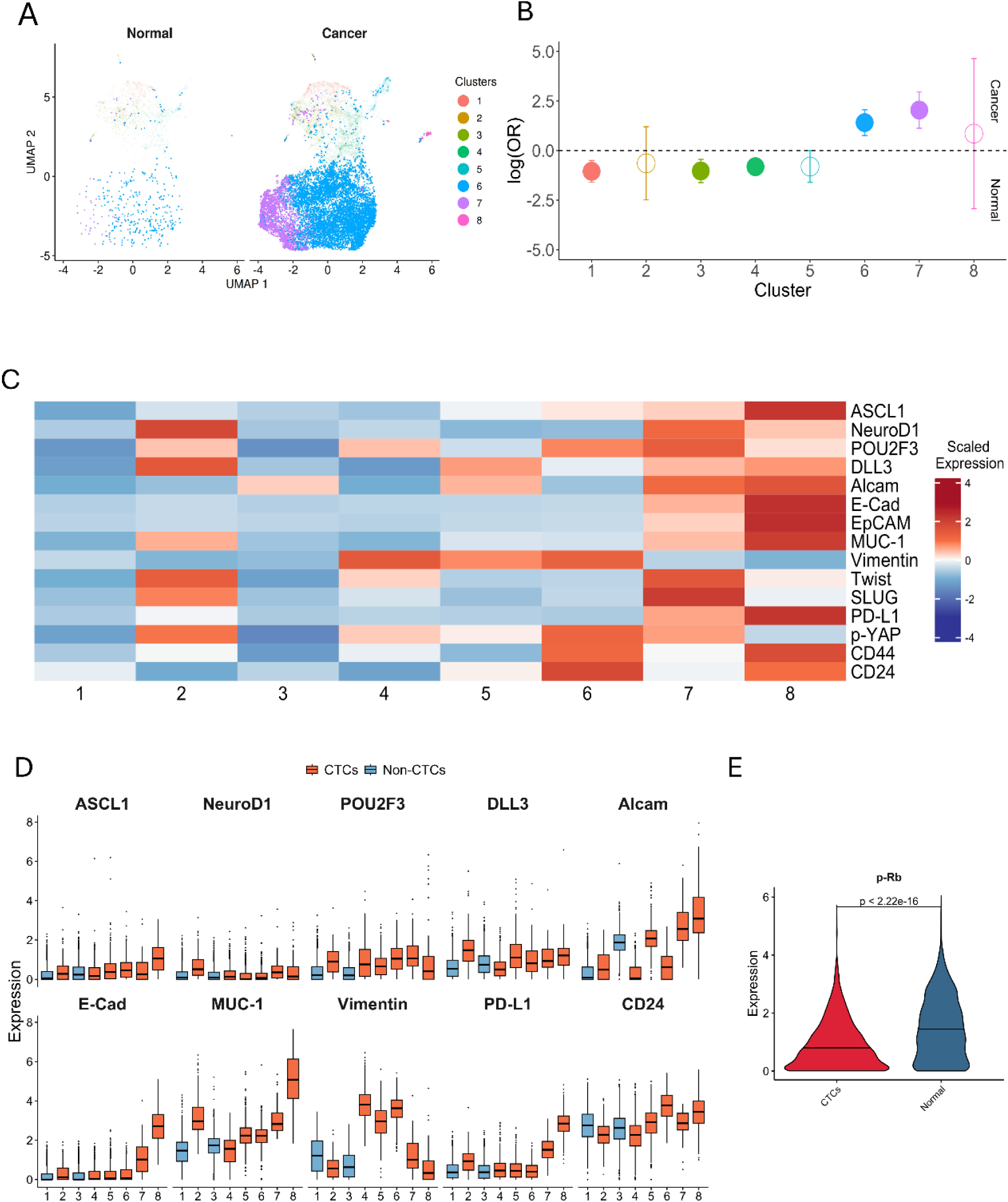
CyTOF-based detection and characterization of CTCs in liquid biopsies. **A.** UMAP visualization of FlowSOM clustering analysis on CD45-CD56+ cells derived from healthy donors (n=1708 cells) vs cancer patients (n=16,255 cells). The three cancer-enriched primary clusters (p6-p8) are shown with opaque colors. **B.** Mixed-effect logistic regression analysis showing the odds ratio (OR) of cluster enrichment in normal vs SCLC liquid biopsies. (P-values for primary clusters p1 to p8: 0.001, 0.9901, 0.0022, 0.0013, 0.1492, 0.0001, 0.0001 and 0.9901, respectively). **C.** Heatmap showing differential marker expression profiles of the eight CD45-CD56+ cell populations resulting from clustering analysis on the cancer-enriched CD45-CD56+ cells identified in panels B and C (Additional details can be found in Supplementary Figure 2). **D.** Boxplots showing the expression levels of markers in the eight new secondary clusters from Figure 1C (clusters s1 and s3 in blue were defined as non-CTC clusters and s2,s4,s5,s6,s7 and s8 in red were designated as CTCs). **E.** Violin plots showing the expression levels of pRB in normal cells vs CTCs. CTCs: CD45-CD56+ cells from SCLC patients that were in clusters s2,s4,s5,s6,s7 and s8 (from C) and Normal cells: all CD45-CD56+ cells detected in healthy donors. (Related to Figure 1B-D). See Methods for additional details.

To refine CTC identification, we performed a secondary clustering analysis on all cells within the three cancer-enriched populations (clusters p6, p7, and p8) which identified eight new distinct cell populations (**Supplementary Figure 2C**) with unique protein expression profiles (**Figure 1C**). Based on elevated (combined or not) expression of SCLC-specific markers (e.g. DLL3), SCLC-subtype-defining transcription factors (ASCL1, NeuroD1, POU2F3) and/or epithelial markers (e.g. EpCAM, E-Cadherin, MUC1) secondary clusters s2 and s4-8 were designated as CTCs, as opposed to clusters s1 and s3 which lacked substantial expression across all markers (**Figure 1C-D**). Moreover, pRB expression was significantly lower in the CTCs compared to “normal cells” (i.e. all CD45-CD56+ cells from healthy donors), corroborating with reports of inactivated or deleted RB protein in SCLC tumors (**Figure 1E**) [19] and further confirming our analysis-driven CTC identification. The average percentage of CTCs per patient in our study was 0.0739% (**Supplementary Table 2**), a number that agrees with recent findings [18, 20, 21]. We did not detect CTCs in 4 out of the 51 patients [MDA-SC355 (LC-NEC), MDA-SC363, MDA-SC370 and MDA-SC400 following our clustering analysis.

### 2.2 Subtype-specific CTC signatures and treatment-associated phenotypic changes

The expression patterns of three lineage-defining transcription factors classify SCLC into four established subtypes: achaete-scute homolog 1 (ASCL1) (SCLC-A), neuronal differentiation 1 (NEUROD1) (SCLC-N), POU class 2 homeobox 3 (POU2F3) (SCLC-P), and the triple negative inflamed (SCLC-I), each linked to unique therapeutic susceptibilities [22]. To evaluate whether further analysis of the CTCs (clusters s2, s4-8 from Figure 1D) would reveal the canonical SCLC subtypes, we analyzed transcription factor expression at a single-cell level within the identified pooled CTC populations. Indeed, clustering identified discrete subpopulations corresponding to the three canonical SCLC molecular subtypes SCLC-A, SCLC-N and SCLC-P (**Figure 2A**), harboring subtype-specific protein expression profiles as previously reported at the transcriptomic level [22] (**Figure 2B**). As expected, neuroendocrine subtypes SCLC-A and SCLC-N showed high expression of DLL3 (an established ASCL-1 transcriptional target [23]), MUC1, E-Cadherin, and EpCAM, substantiating their epithelial nature whereas SCLC-P exhibited higher expression of mesenchymal markers, namely, Vimentin, and lower expression of neuroendocrine markers consistent with previously reported findings (**Figure 2B**) [24, 25]. Finally, our analysis revealed an ASCL1/NeuroD1/POU2F3 low, mesenchymal group of cells (high vimentin/absence of epithelial markers E-Cadherin, EpCAM and MUC1) and defined it as the SCLC-Mes subtype.

**Figure 2.**
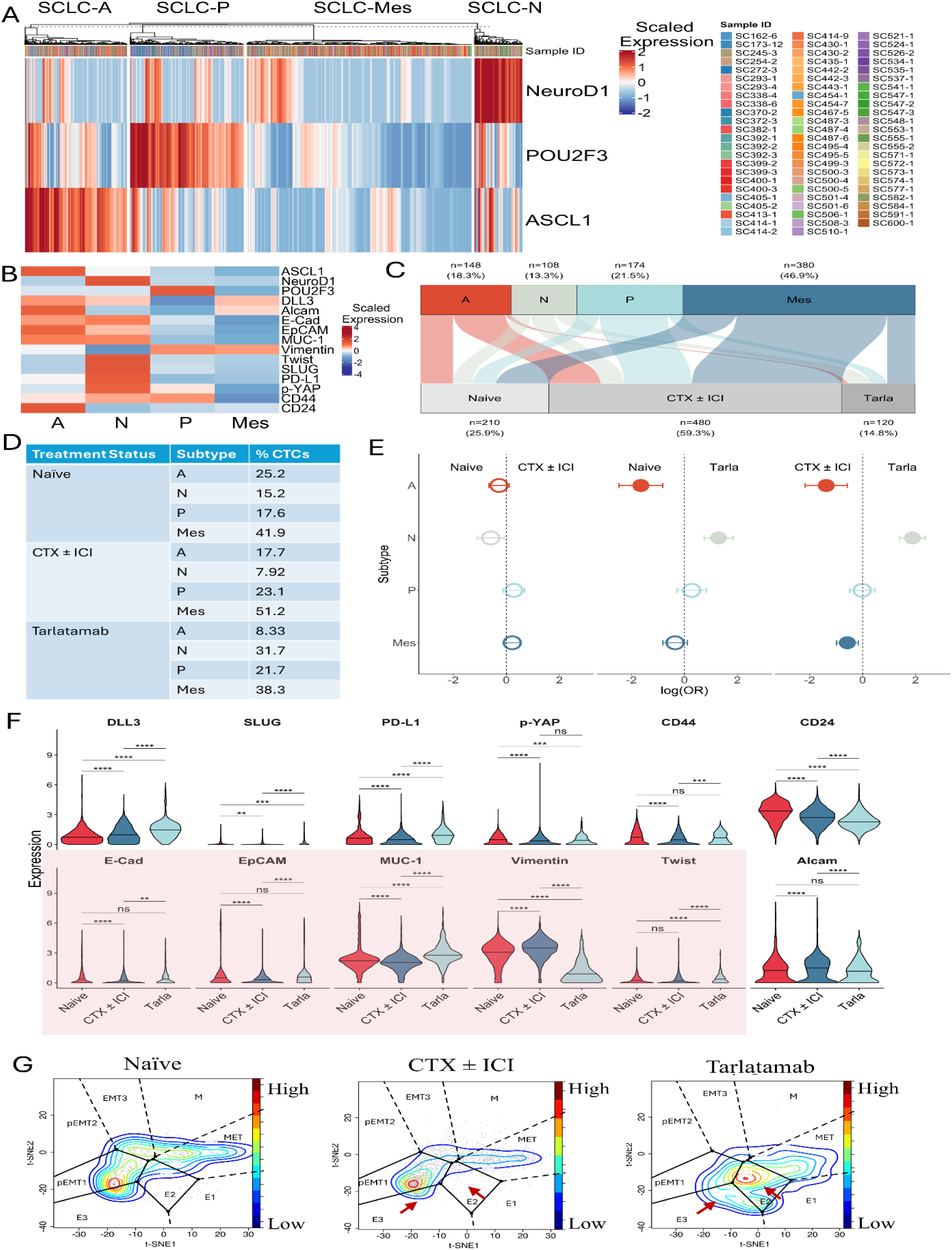
Identification of canonical SCLC molecular subtypes and associated protein profiles in CTCs detected in liquid biopsies. **A.** Dendrogram showing the three established SCLC subtypes, SCLC-A, SCLC-P, SCLC-N and SCLC-Mes subtype identified in CTCs (Total #cells: 3291) from SCLC patient liquid biopsies. Clustering (k-means clustering (k=4)) was performed based on protein expression of transcription factors ASCL1, NeuroD1 and POU2F3. Top annotations indicate the sample (patient) ID from which each CTC was derived. **B.** Heatmap showing protein marker expression profile of CTCs within each subtype. **C.** Alluvial plot showing the distribution of CTCs across treatment groups after equal down-sampling to 30 cells per liquid biopsy (28 liquid biopsies with at least 30 CTCs/specimen from 23 patients). Each flow represents the proportion of cells assigned to a subtype within each treatment category. **D.** Table summarizing the percent of CTCs of each subtype in each treatment group as shown in the alluvial plot in C. **E.** Odds ratio (OR) forest plots displaying associations between CTC subtypes and treatment groups (left; naïve vs CTX±ICI, middle; naive vs tarlatamab, right; CTX±ICI vs tarlatamab), calculated using the chi-squared test. Log(OR) > 1 indicates enrichment of a subtype in the right treatment condition relative to all other subtypes, while log(OR) < 1 indicates enrichment of a subtype in the left treatment condition relative to all other subtypes. (Naive vs CTX±ICI P values: A: 0.8007, N: 0.3145, P: 0.8007, I: 0.8007) (Naive vs tarlatamab P Values: A:_2.269e-04, N: 1.883e-05, P: 8.007e-01, I: 8.007e-01) (CTX±ICI vs tarlatamab P values: A: 3.122e-03, N: 4.378e-14, P: 9.175e-01, I: 6.557e-02. Additional information provided in Supplementary Figure 3. **F.** Violin plots showing differential marker expression in all CTCs from naïve vs CTX±ICI vs tarlatamab-treated samples. Shaded areas indicate EMT-related markers that drive PHENOSTAMP projections in G. **G.** PHENOSTAMP projections showing the enrichment of CTCs at a single-cell level in different EMT states from treatment naïve (left; n=1038), CTX±ICI treated (middle; n=2013) and tarlatamab-treated patients (right; n=240). Red arrows indicate where the most prominent changes in EMT state enrichment were observed. Red color indicates higher cell density and blue color indicates lower cell density.

Importantly, this subtype pattern and marker expression profile was not recapitulated when the same analysis was performed on CD45-CD56+ cells in the healthy blood enriched primary clusters p1-p5 from Figure 1B or from the non-CTC secondary clusters s1 and s3 from Figure 1D, confirming our CTC-designated cell clusters (**Supplementary Figure 3A**). When we interrogated subtype distribution of CTCs at the liquid biopsy/patient level, we observed inter-as well as intra-patient CTC subtype heterogeneity in liquid biopsies with at least 10 CTCs detected **(Supplementary Figure 3B).** When available, we validated our CyTOF/CTC-based findings by comparing them to proteomic or single-cell RNA sequencing data from matched primary tumor tissue or patient-CTC derived xenografts, enabling subtype concordance between primary tumor and CTCs (**Supplementary Figure 3C and D**).

Next, we investigated subtype association with treatment modality in CTCs from treatment naïve vs. patients undergoing therapy, which includes platinum-based chemotherapy alone or with immunotherapy (CTX ± ICI) as well as the new anti-DLL3 therapy. Tarlatamab, a DLL3-targeted BiTE, was recently approved by the FDA (2024) for treatment of extensive stage-SCLC, marking a new therapeutic avenue for relapsed SCLC [26]. Most patients receiving tarlatamab therapy in our cohort had previously undergone platinum-based chemotherapy and anti-PD-L1 treatment (CTX±ICI). To assess treatment-associated subtype enrichment in CTCs and avoid potential overrepresentation of specimen-specific CTCs, we interrogated CTCs from equally down-sampled specimens (30 CTCs per liquid biopsy). This resulted in assessing subtype association with treatment modality in 810 CTCs from 27 liquid biopsies/samples originating from 21 patients that met the criterion of at least 30 CTCs/sample (210 CTCs from 7 naïve samples from 7 patients, 480 CTCs from 16 CTX±ICI-treated samples from 13 patients and 120 CTCs from 4 tarlatamab-treated samples from 3 patients) (**Supplementary Figure 3B, Supplementary Table 3**).

Figure 2C shows a single-cell alluvial plot of CTC subtype distribution across treatment naïve, CTX±ICI treated, and tarlatamab groups. In our cohort of SCLC liquid biopsies, CTC subtype distribution was dominated by SCLC-Mes (46.9%), followed by SCLC-P (21.5%), SCLC-A (18.3%), and SCLC-N (13.3%). In contrast, primary tissue datasets report approximate proportions of 35% SCLC-A, 35% SCLC-N, 15% SCLC-P, and 15% inflamed (SCLC-I) subtypes [27]. This divergence may reflect both biological differences between primary tumors and CTCs and the fact that our cohort predominantly includes treated patients known to undergo a shift toward non-neuroendocrine, more mesenchymal phenotypes [22]. Moreover, SCLC-Mes was also the most prevalent subtype within each treatment modality: 41.9%, 51.2%, and 38.3% in naïve, CTX±ICI, and tarlatamab specimens respectively. SCLC-A, SCLC-N and SCLC-P CTCs were most enriched in naïve (25.2%), tarlatamab (31.7%), and CTX ± ICI (23.1%) specimens respectively (Figure 2D).

Given the relatively small number of CTCs analyzed, we performed a chi-square test to assess statistical significance of subtype associations with treatment setting (Figure 2E). The distributions of subtype proportions across 1000 iterations of random sampling of 30 CTCs per liquid biopsy are shown in **Supplementary** Figure 3E. Our analysis showed associations of specific subtypes with each of the three treatment modalities, albeit not all reaching statistical significance. Specifically, when comparing the treatment naïve setting with CTX±ICI and tarlatamab settings, the former was associated with subtype SCLC-A, a feature established in naïve primary biopsy specimens [16].CTX±ICI treatment was more associated with subtype SCLC-P and to a less degree with SCLC-Mes, consistent with previous findings [22–24]. Tarlatamab treatment was significantly associated with an enrichment of SCLC-N CTCs. SCLC-N enrichment in the tarlatamab setting was also observed when compared to CTX±ICI (most of the tarlatamab treated patients in our cohort were previously on CTX±ICI) (Figure 2E). Importantly, our findings were comparable to when we analyzed all CTCs detected across our entire cohort of liquid biopsies (without down-sampling), specifically in terms of relative proportions of each subtype within each treatment modality and associations among subtypes and treatment settings, especially in the cases between CTX±ICI with SCLC-P and tarlatamab with SCLC-N, where we applied a linear mixed-effects regression model to account for inter-patient variability (alluvial plot, table of subtype distributions, and statistical analysis of all CTCs provided in **Supplementary** Figures 3F-H).

In addition to assessing CTC subtype association with treatment modality, we also investigated differences in all other markers of our CyTOF panel, including EMT-related markers (Figures 2F, and **2G**). When comparing CTCs from CTX±ICI and tarlatamab-treated patients, the most notable differences were observed in PD-L1, CD24 and DLL3 expression. Under CTX±ICI conditions, PD-L1 and CD24 decreased when compared to treatment naïve conditions, whereas in tarlatamab-treated conditions PD-L1 was increased, and CD24 further decreased compared to both naïve and CTX±ICI conditions. Interestingly, DLL3 significantly increased in both treatment modalities compared to the naïve setting, with tarlatamab CTCs exhibiting a more notable increase in DLL3 when compared to both naïve and CTX±ICI - treated CTCs.

EMT has been shown to drive both intrinsic and acquired resistance in SCLC, especially to chemotherapy, by enhancing cellular plasticity and upregulation of pro-survival pathways [28, 29]. Therefore, we interrogated EMT changes in relation to treatment. Under CTX±ICI, CTCs showed a moderate increase in Vimentin levels and decreased levels of E-Cadherin, EpCAM and MUC1 (Figures 2F), suggesting a shift toward a more partial EMT (pEMT) post-treatment. In contrast, tarlatamab-treated CTCs exhibited significantly reduced Vimentin expression alongside increased levels of E-Cadherin, EpCAM, and MUC1 compared to both naïve and CTX±ICI - treated CTCs, indicating induction of an epithelial phenotype (Figure 2F, pink shaded area). Our EMT observations were confirmed when we projected CTCs from all three conditions on PHENOSTAMP, a lung cancer reference map of EMT and MET (mesenchymal-to-epithelial transition) states referred to as the EMT-MET PHENOtypic STAte MaP, previously developed by our group [14, 30]. Specifically in the case of CTX±ICI, we observed a minor enrichment of a pEMT state with concomitant decrease of epithelial states (E1, E2), whereas projections of tarlatamab CTCs revealed a marked enrichment of all three epithelial states when compared to both naïve and CTX±ICI - treated CTCs (Figure 2G).

### 2.3 Temporal tracking of individual patients reveals treatment-associated changes in subtype proportions and biomarker expression in CTCs

We next wanted to explore the functional implications of phenotyping liquid biopsy CTCs in *in vivo* xenograft models from which we had matching liquid biopsy CyTOF data. Figure 3A shows ASCL1 expression in xenograft models [one patient-derived xenograft (PDX) and two CTC-derived xenografts (CDXs) respectively] established from three treatment-naïve SCLC patients: one from a primary tumor biopsy (MDA-SC293) and two from pleural effusion CTCs (MDA-SC506 and MDA-SC443). CTCs from the blood of these three patients displayed distinct subtype compositions, with MDA-SC293, MDA-SC506, and MDA-SC443 showing progressively higher ASCL1 expression and neuroendocrine features in that order. MDA-SC443 specifically harbored >50% ASCL1+ CTCs, which was confirmed by immunohistochemistry (IHC) on the respective xenograft tumor (Figure 3B). Although all three models were derived from treatment-naïve patients, they exhibited distinct tumor growth kinetics, and increased latency periods that positively correlated with elevated ASCL1 expression in both CTCs and the tissue of origin (Figure 3C), which, in turn, also correlated with a predominantly epithelial phenotype as confirmed by our PHENOSTAMP projections (Figure 3D **and Supplementary** Figure 4A). To further support this link, we demonstrated concordant protein expression profiles in CD45-CD56+ cells detected in both a pleural effusion and blood sample acquired from patient MDA-SC443 (**Supplementary** Figure 4B).

**Figure 3.**
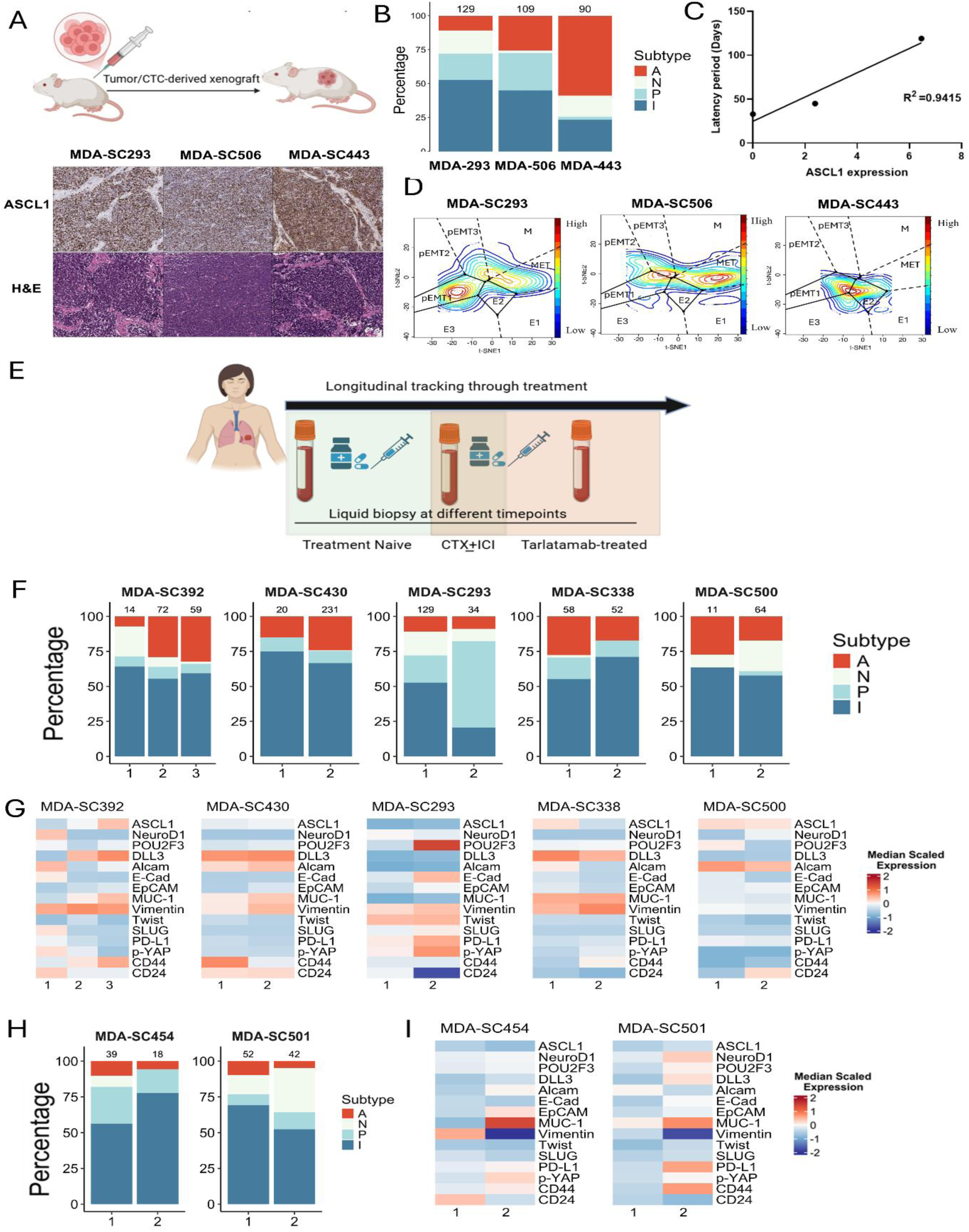
Temporal tracking of individual patients reveals treatment-associated subtype transitions and changes in CTC protein markers A. (Top) Schematic illustration showing subcutaneous injection of patient-derived xenografts in mice and (bottom) immunohistochemistry images showing the expression of ASCL1 in the three patient xenograft tumors derived from liquid biopsies of patients MDA-SC293, MDA-SC506 and MDA-SC443. B. Stacked bar graphs showing the CTC subtype composition from our CyTOF analysis of each of the three naïve patient liquid biopsies from A. Numbers on top of each bar indicates number of CTCs as defined by our computational pipeline. C. Linear regression graph showing correlation of median ASCL1 expression in CTCs to latency period (number of days passed before a palpable tumor was formed/detected) in each patient-derived xenograft. D. PHENOSTAMP showing the enrichment of CTCs in different EMT states from patients MDA-SC293, MDA-SC506 and MDA-SC443 (Note: The patient of which majority of CTCs were SCLC-A was in fact more epithelial). Red color indicates higher cell density and blue color indicates lower cell density. Additional information provided in Supplementary Figure 4). E. Schematic illustration showing the longitudinal tracking of patients before and after therapy. F. Stacked bar plots showing the percentage of CTCs in each subtype within a patient at different timepoints during CTX±ICI (MDA-SC293-1 was a treatment-naïve sample). G. Heatmaps showing median scaled expression for each marker in CTCs from each patient at different timepoints. H. Stacked bar plots showing the percentage of CTCs in each subtype within a patient at different timepoints during tarlatamab therapy (MDA-SC454-1 was a treatment-naïve sample). I. Heatmaps showing median scaled expression for each marker in CTCs from each patient at different timepoints during tarlatamab therapy. The number underneath the heatmaps denotes the n^th^ number of blood draw or time point of the liquid biopsy from that patient. The number on top of each stacked bar denotes the number of CTCs as defined per our computational pipeline (See Supplementary Figure 2A).

To explore dynamic CTC phenotypes and treatment-related changes in subtype proportions at the individual level, we profiled CTCs from seven patients before and after treatment (Figure 3E). For personalized analysis of marker expression, we included patients with ≥10 CTCs detected per timepoint. Five of eight CTX±ICI - treated and two of seven tarlatamab-treated patients met this criterion (Figure 3F**–I**). Two CTX±ICI - treated patients (MDA-SC392 and MDA-SC430) that showed stable disease at latest collection, had increased proportion of neuroendocrine CTCs and ASCL1 expression over the course of therapy, along with a progressive decrease in p-YAP (Y357) and PD-L1 levels (Figure 3F and G). These observations are consistent with prior reports linking high YAP1 and PD-L1 expression to poorer survival outcomes and more advanced disease in SCLC patients [31]. Conversely, the three other CTX±ICI treated patients (MDA-SC293, MDA-SC338 and MDA-SC500) who experienced disease progression displayed a reduced proportion of neuroendocrine SCLC-A CTCs, with patients MDA-SC293 and MDA-SC338 also showing concomitant increase in SCLC-P and SCLC-mes subtype proportions and upregulation of p-YAP, and PD-L1 levels, further supporting the relevance of CTC phenotyping in tracking disease dynamics (Figure 3F and G).

In the two patient cases who had transitioned from CTX±ICI to tarlatamab treatment and had experienced disease progression, post-tarlatamab CTCs exhibited different subtype proportions albeit both showing a small decrease in SCLC-A CTC proportion with time (Figure 3H). Moreover, in both cases, post-tarlatamab CTCs expressed higher MUC1, PD-L1, p-YAP and DLL3 levels and significantly lower Vimentin levels compared to pre-tarlatamab CTCs (Figure 3I). These findings underscore the value of our multiplexed approach, which enables detection and interrogation of CTC-specific subtypes, phenotypic markers and therapeutic targets (e.g. DLL3) from liquid biopsies collected in real time before and after diverse treatment regimens.

## 3. Discussion

We present the first high-dimensional single-cell proteomic analysis of CTCs detected directly in SCLC liquid biopsies, revealing canonical molecular SCLC subtypes in circulation using a minimally invasive approach that enables real-time monitoring. Our approach, which combines targeted marker-driven gating with phenotypic analysis of tumor and lineage-associated proteins, represents a significant advancement over traditional CTC detection methods like CellSearch, which relies on a single marker such as EpCAM-known to fluctuate with EMT, also shown in our UMAP analysis to be expressed in only ∼30% of SCLC cell lines (Figure 1 **and Supplementary** Figure 1B) [18], and Parsortix, which cannot detect small diameter cells [32]. A recent study demonstrated that CyTOF provides greater resolution and depth than bulk RNA sequencing in characterizing CTC subpopulations and immune phenotypes in Head and Neck Squamous Cell Carcinoma (HNSCC) [33]. Unlike that study, which used Parsortix-based CTC enrichment, our approach enables direct detection of CTCs and their current phenotypes in liquid biopsies without enrichment, avoiding phenotype alterations related to *in vitro* cell culture conditions. By incorporating both surface and intracellular markers, we achieved *in silico* enrichment of tumor-derived cells and captured broader SCLC heterogeneity, highlighting the enhanced sensitivity and practicality of our method. Our CyTOF approach allowed us to phenotypically resolve distinct CTC populations corresponding to the SCLC-A, SCLC-N, and SCLC-P subtypes, long established at a transcriptomic level in tissue but previously uncharacterized in CTCs and also helped identify a SCLC subtype characterized by absence/low expression of NeuroD1, ASCL1 and POU2F3 and presence of mesenchymal features (SCLC-mes). Although in our analysis this subtype was not high in p-YAP expression, we cannot exclude the possibility that this subtype overlaps/includes subpopulations of cells of the previously described inflammatory SCLC-I/Y/H subtypes [34, 35]. Future studies necessitate incorporation of inflammatory markers in our SCLC CyTOF panel to further characterize the SCLC-mes subtype and test whether it in fact overlaps with the SCLC-I subtype previously reported by our group [22].

A key finding of our study is the detection of dynamic changes of CTC subtype proportions in patients with progressive or relapsed disease (Figure 3) - plasticity not captured by cell-free DNA-based methods [11, 36]. While such transitions have been postulated in tumor tissue and preclinical models, our data provide the first real-time, minimally invasive evidence of changes in subtype proportions in circulation. Given the growing interest in subtype-specific vulnerabilities in SCLC [22], these insights have important clinical implications for therapeutic stratification and early resistance detection. We observed a general shift of CTC proportions from neuroendocrine to non-neuroendocrine subtypes after treatment, but variable patient-specific patterns revealed by longitudinal single-cell profiling underscore the importance of temporal sampling to guide precision therapy. Although the number of longitudinally sampled patients in this study was modest, our data illustrate the limitations of single time-point primary tumor biopsies in capturing the temporal dynamics of a highly plastic disease such as SCLC.

Interestingly, our findings highlight that higher ASCL1 expression observed both in CDX models with longer tumor latency and in patients with stable disease, defines an epithelial phenotype that may be associated with less aggressive CTCs. These patients also had reduced levels of p-YAP and PD-L1 post therapy. In contrast, patients with declining ASCL1 expression exhibited elevated levels of p-YAP and PD-L1 and transitioned toward mesenchymal phenotypes following CTX±ICI. (Figure 3G). Reportedly, phosphorylation of YAP at Tyrosine 357 is an activating modification that helps in translocation of YAP into the nucleus, facilitating its activity as a transcriptional coactivator [37]. These findings are consistent with previous reports that associate elevated PD-L1 expression, regulated by YAP1, with worse survival outcomes in a specific subtype and more advanced stages of SCLC [31] and support the hypothesis that CTC phenotypes are not static, and that phenotypic plasticity-especially in response to therapy-may underlie mechanisms of resistance in SCLC.

Importantly, we provide the first single-cell proteomic CTC data from patients treated with tarlatamab based on samples collected between 2-12 months after treatment initiation. In these patients, we observed dynamic shifts in both marker expression and CTC subtype composition, offering mechanistic insights into treatment adaptation and probable onset of early resistance. Interestingly, while patients receiving CTX±ICI exhibited a progressive, modest enrichment of pEMT and mesenchymal features-consistent with selection for more invasive, SCLC-P-like phenotypes; those who received tarlatamab displayed an increased enrichment of neuroendocrine, epithelial states and decreased mesenchymal traits, more representative of the SCLC-N subtype (Figure 2E-G). This suggests that targeted treatment and immunotherapy may select for or induce phenotypically plastic cell states, including EMT-associated transitions, which are often linked to therapy resistance and metastatic potential. Although DLL3 is a neuroendocrine marker, the observed pro-neuroendocrine shift in post-tarlatamab CTCs suggests therapy-induced reprogramming or selective elimination of mesenchymal-like cells, implying DLL3-targeted therapy may influence tumor plasticity beyond cytotoxicity. Whether tarlatamab promotes the selection of DLL3-low, plastic cells that could later re-express DLL3 remains to be explored. However, a prior study in neuroendocrine prostate cancer has shown that following T-cell mediated elimination of DLL3+ cells, residual DLL3-low tumor cells can persist and exhibit phenotypic plasticity, with some relapsed tumors re-expressing DLL3, suggesting dynamic and reversible regulation of DLL3 under therapeutic pressure [38]. This indicates that our approach could potentially help stratify patients for targeted therapies, for instance, anti-DLL3 BiTEs since we can detect DLL3 levels in circulation. The upregulation of PD-L1 and p-YAP in CTCs from tarlatamab-treated patients (including the ones tracked longitudinally) (Figures 2F**, 3I**) suggests that some tumors may undergo phenotypic changes associated with adaptive immune evasion and emerging resistance to immunotherapy, which could have important implications for subsequent treatment responsiveness and influence survival outcome in the clinic. MUC1, a marker known to be associated with increased oncogenic potential, drug resistance, anoikis-resistance and MET in epithelial cancers including SCLC [14, 15, 39], was also found to be upregulated in tarlatamab-treated patients (Figures 2F, **3H and 3I**). MUC1 has been shown to promote self-renewal and tumorigenicity and maintain neuroendocrine features in SCLC [40, 41], suggesting its potential role in conferring resistance to DLL3-targeted therapies by supporting tumor cell survival and plasticity. However, the precise roles of MUC1, PD-L1 or p-YAP in this case remain to be elucidated and warrants further investigation. By longitudinally profiling CTCs in patients receiving CTX±ICI and tarlatamab therapies, we demonstrate that treatment-induced phenotypic shifts-can be noninvasively captured and may function as early indicators of resistance onset. These observations suggest that co-targeting YAP/PD-L1 signaling, or MUC1-mediated pathways could potentiate tarlatamab activity and delay or prevent future therapeutic resistance.

### Limitations

Despite the strengths of our approach, several limitations remain. The number of healthy donor controls and patient samples may impact the specificity of tumor versus non-tumor classification. Additionally, although longitudinal sampling was included, the relatively few time-points and low CTC counts-especially in the tarlatamab-treated cohort-limit the generalizability of our findings across the full disease course. To address these constraints, we are actively expanding our patient cohort and clinical datasets. We are also in the process of integrating genomic and transcriptomic data for a more comprehensive analysis of the identified CTC populations. Future work will also incorporate immune profiling of patient samples to explore interactions and associations between CTCs and immune cells. This is particularly important in the context of immunotherapy, where identifying correlations between tumor and immune cell states may uncover predictive biomarkers of treatment response or resistance and help stratify patients for treatment. Overall, our study provides a clinically relevant framework for high-resolution, real-time phenotypic profiling of SCLC CTCs through liquid biopsies. By capturing molecular subtypes, monitoring tumor plasticity, and assessing treatment-driven CTC changes, we demonstrate the potential of CyTOF-based single-cell profiling to inform adaptive therapeutic strategies in SCLC.

## Resource availability Lead contact

Further information and requests for resources and reagents should be directed to and will be fulfilled by the lead contact, Loukia G. Karacosta.

## Materials availability

This study did not generate any new unique reagents.

## Data and code availability

All newly generated CyTOF data output files, code generated and utilized for analyses, are provided on github (https://github.com/coleruoff/sclc_cytof). Any additional information required to analyze data will be made available upon request to the lead contact.

## Acknowledgements

This work was supported by DOD/CDRMP HT9425241070 and MDACC research start-up funds (L.G.K.), University of Texas SPORE in Lung Cancer (NIH/NCI P50-CA070907) (L.G.K. and L.A.B.), and NIH/NCI R01-CA207295, NIH/NCI U01-CA256780, MD Anderson Cancer Center CCSG (NIH/NCI P30-CA016672), NIH/NCI R50-CA243698, CPRIT RP210159, generous philanthropic contributions to The University of Texas MD Anderson Cancer Center Lung Cancer Moonshot Program and The Rexanna Foundation for Fighting Lung Cancer (L.A.B.). We would like to thank the Flow Cytometry and Cellular Imaging Core Facility that was supported in part by The University of Texas MD Anderson Cancer Center and P30CA016672. We would also especially like to thank A.R.K., L.W.Y., M.J.A., R.B.N., K.E.N., J.O., J.K.R., B. & B.N, C.K., P.C.B., S.S., S.R., and W.A.B. for their philanthropic support of these projects.

## Author contributions

Conceptualization, L.G.K, M.B., and C.R.; data curation, M.B., C.R., S.E., C.A.S., T.W., A.D., R.W., B.Z.; formal analysis, M.B., C.R., S.E., C.A.S., J.L., B.A., L.D., J.W., L.S.S., A.G.S. and L.G.K.; funding acquisition, L.A.B and L.G.K; investigation, M.B., C.R., S.E., C.A.S., S.B., L.S.S., A.G.S. and L.G.K.; methodology, M.B., C.R., S.E., C.A.S., S.B., T.W., F.E.L., A.V. and L.G.K.; project administration, C.A.S., A.D., A.V., R.C.; software, C.R., S.E., M.B., J.L., B.A. and L.G.K.; resources, C.M.G, L.A.B. and L.G.K; supervision, L.A.B. and L.G.K, validation, M.B., C.R., S.E., C.A.S., F.E.L., R.W., B.Z., C.M.G., L.G.K.; visualization, M.B., C.R., S.E., J.L., B.A., L.G.K.; writing – original draft, M.B., C.R. and L.G.K; writing – review & editing, all authors reviewed the final manuscript.

## 4. Star Methods

### 4.1 Cell Culture

SCLC cell lines H1105, H526, H196, H209 and NJH29 were provided from Dr. Lauren Byers at MD Anderson and were grown in RPMI-1640 and DMEM (Dulbecco’s modified Eagle’s medium) media respectively, supplemented with 10% fetal bovine serum (FBS), and 5% antibiotic solution (penicillin/streptomycin), at 5% CO_2_ and 37 °C. In certain runs, SCLC cell lines purchased from ATCC (Manassas, VA) were used. NJH29 cells were originally generated in Julien Sage’s lab at Stanford University [42].

### 4.2 Western blotting, immunohistochemistry, and single-cell RNA sequencing

Generation of patient tumor-derived or CTC-derived xenografts, western blotting, immunohistochemistry and sc-RNA sequencing were performed on the primary tumors or cell-derived xenograft tumors using previously published protocols [43].

### 4.3 Study cohorts, sample collection and sample processing

This study was conducted in accordance with the provisions of the Declaration of Helsinki and Good Clinical Practice guidelines. The project was performed under The University of Texas, MD Anderson Cancer Center Institutional Review Board approved protocol (PA13-0589, PA14-0531, LAB10-0442) with informed consents. Blood samples were obtained from 19 healthy donors, 18 treatment-naïve (18 liquid biopsies) and 31 (SOC) treated patients (43 liquid biopsies collected across different time points), and 24 of those samples were from 10 longitudinally tracked SCLC patients. We also received 14 pre-tarlatamab (14 patients) and 14 post-tarlatamab liquid biopsies from 10 patients undergoing tarlatamab therapy (4 longitudinally tracked biopsies). Venous blood was sampled directly into a 10 ml EDTA or heparin-blood collection tube (BCT). Whole blood was used to isolate peripheral blood mononuclear cells (PBMCs) using Ficoll-Paque plus according to a previously described protocol [15]. The workflow is shown in Figure 1A and the number of patients and CTCs used for each computational analysis step and type is shown in Table 1 and **Supplementary Table 3**.

### 4.4 Liquid biopsy processing for CyTOF analysis

Fresh blood samples from healthy donors were obtained either from the blood bank or consented volunteers at MDACC and placed in 4°C until processing on a rotator. Following PBMC isolation, a hemocytometer or automated cell counter (Trypan Blue exclusion) were used for counting and initial % viability assessment of PBMCs which were resuspended in phosphate-buffered saline (PBS). In the few cases where blood specimens were left in 4 °C overnight, no significant effect was observed in viability as assessed with Trypan Blue counting. For assessing cell viability with CyTOF, cell pellets were subsequently incubated in 1mL PBS containing Cell ID Cisplatin (Fluidigm, final concentration 2.5 μM) for 5 min at room temperature. The reaction was quenched by adding an equal volume of complete media containing 10% fetal bovine serum (FBS), and cells were immediately washed twice (centrifuged for 5 min at 500×g) with complete media to remove excess cisplatin. Cell pellets were resuspended in complete RPMI media to reach 0.5–1 × 10^6^ cells/mL aliquots per sample and fixed by adding paraformaldehyde (PFA) at a final concentration of 1.6% for 10 min at room temperature. Fixed cells were centrifuged and washed twice at 500×g for 5 min at 4 °C to remove the residual PFA with Cell Staining Media (CSM, 0.5% w/v BSA, 0.02% w/v NaN3 in PBS). Processed cells were cryopreserved at −80°C in CSM until further use in downstream CyTOF staining and analysis.

### 4.5 CyTOF antibodies

Details on antibodies used in the CyTOF panel, including information on antibody clone, vendor, species reactivity, catalog number and conjugated metal isotope are summarized in **Supplementary Table 1**. Antibodies were either purchased/provided by the MD Anderson CyTOF Core Facility or conjugated to metal isotopes in-house using the MaxPar Antibody Conjugation Kit (Fluidigm) and titrated to determine optimal staining concentrations as described previously [15].

### 4.6 Antibody staining procedure and CyTOF analysis

Previously fixed and cryopreserved PBMCs were retrieved from −80°C storage and thawed on ice. Based on cell counts obtained prior to freezing, sample volumes corresponding to 1 million cells were calculated. Thawed cells were washed twice with CSM to remove residual fixative and then transferred into individual 2 mL cluster tubes.

Cells were resuspended in 800 µL of 1× Barcode Perm Buffer (Fluidigm), and individual barcodes from the palladium-based Cell-ID™ 20-Plex Pd Barcoding Kit (Fluidigm) were reconstituted in 100 µL of Barcode Perm Buffer and mixed thoroughly. For each sample, 100 µL of the appropriate barcode was added to the corresponding tube. Samples were mixed gently and incubated for 30 minutes at room temperature to allow for barcoding. After incubation, cells were washed twice with CSM to remove excess barcode reagents.

All barcoded samples were then pooled into a single 5 mL round-bottom FACS tube and washed once more with CSM. The pooled cells were stained for surface marker expression by adding surface antibody cocktail (per core instructions) provided by the MD Anderson Flow Cytometry Core Facility and surface antibodies conjugated in-house. The staining mixture was incubated for 30 minutes at room temperature. Following surface staining, cells were washed once with CSM and permeabilized by adding ice-cold methanol, followed by incubation on ice for 10 minutes. Methanol was then washed off with CSM. Intracellular staining was performed by adding intracellular antibody cocktail (per core instructions) and intracellular antibodies conjugated in-house and incubating the mixture for 30 minutes at room temperature (**Supplementary Table 1**). After staining, cells were washed again with CSM. For DNA labeling, cells were resuspended in intercalation solution consisting of 1 part 16% paraformaldehyde in 9 parts PBS, supplemented with 1:5000 dilution of Cell-ID™ Intercalator-Ir (Fluidigm). The cells were incubated in this solution for 20 minutes at room temperature and stored overnight at 4°C. On the following day, samples were washed once with CSM, followed by two washes with ultrapure water. Prepared samples were then acquired on a Helios™ mass cytometer (Fluidigm) according to the manufacturer’s standard operating procedures at the core facility.

### 4.7 Processing CyTOF Data

Data from each sample were de-barcoded and batch normalized as described previously [44]. Using Cytobank, we excluded beads, dead cells, doublets, and apoptotic cells using manual gating (Supplementary Figure 1A). A final gating was done on CD45-CD56+ cell populations. Individual untransformed ‘.fcs’ files for each sample were read into R using read.flowSet() from the flowCore package (v2.18.0). The flowset was then converted to a SingleCellExperiment (v1.28.1) object using prepData() from the CATALYST package (v1.30.2). The data was subsequently transformed using the arcsinh function with a cofactor of 5.

### 4.8 Quality Control

CyTOF runs that were determined to be clear outliers by individual protein marker visualization were removed from the analysis. Samples with low viability (< 75% viable cells by Cisplatin staining) were also removed from the analysis. Because most samples were run more than once for technical replicability assessment, we performed proportional down-sampling on all cells from each liquid biopsy sample proportional to the number of times the sample was run on CyTOF. [percent sampled = 1/(#of runs)].

For individual CyTOF runs we included internal controls (reference SCLC cell lines NJH29, H526, and H1105 representative of SCLC subtypes - N, P and A respectively) per batch, for evaluating SCLC specific marker expression. Blood specimens from healthy donors were also included in batches as negative CTC controls. Principal component analysis (PCA) confirmed minimal batch effects across all CyTOF runs, with consistent trends observed for replicate samples across multiple runs and repeat CyTOF runs demonstrated high reproducibility of marker expression trends over internal control marker expression, validating technical replicability (**Supplementary** Figures 1D**, E and F**).

### 4.9 Clustering and Dimensionality Reduction

To identify cellular subpopulations within the data, we applied the FlowSOM clustering algorithm using the default parameters of the cluster() function from the CATALYST package. We then merged the initial clustering results into 8 consensus clusters. Consensus clustering was performed using the ConsensusClusterPlus R package (v1.70.0) to assess the stability of unsupervised clustering results across resampled datasets. The optimal number of clusters (k) was selected by examining the cumulative distribution function (CDF) and delta area plots. The delta area plot, which quantifies the increase in consensus (stability) achieved when increasing k, showed that improvements in cluster stability plateaued beyond k = 8 (**Supplementary** Figure 2B). Therefore, 8 clusters were selected as the most stable and biologically relevant solution.. Dimensionality reduction (UMAP) was utilized to visualize distinct subpopulations but did not influence the clustering methodology. SCLC cell lines and CD45⁻ cells from patient samples clustered within the same high-dimensional space marked by high CD56 expression, supporting the use of CD45⁻CD56⁺ gating to identify CTCs (Supplementary Figure 1B).

To exclude non-CTC-like cell populations, we applied a two-step unsupervised clustering strategy (**Supplementary** Figure 2A). In the first step, we performed clustering using the FlowSOM algorithm on all pooled CD45⁻CD56⁺ cells derived from 19 healthy donor blood samples and 75 liquid biopsies from 51 patients with SCLC (Figure 1B; **Table 1; Supplementary Table 3**). The source of each individual CD45-CD56+ cell was not fed into the algorithm prior to applying FlowSOM, and then this unsupervised clustering analysis identified eight distinct clusters namely primary clusters p1-p8. Among these, clusters p6, p7, and p8 were more enriched in samples from SCLC patients than in those from healthy donors, with clusters p6 and p7 showing statistically significant enrichment. Based on this pattern, clusters p6-p8 were collectively defined as cancer-enriched clusters (Figure 1C). Although cluster p8 did not reach statistical significance, it was included due to its exclusive presence in a single patient sample. UMAP was utilized to visualize distinct subpopulations but did not influence the clustering methodology.

In the second step, we further analyzed all cells within the cancer-enriched clusters (p6-p8) using additional unsupervised clustering, which revealed eight new subpopulations with distinct protein expression profiles (**Supplementary** Figure 2B and C; Figure 1D). Clusters s2, s4, s5, s6, s7, and s8 were classified as CTCs based on elevated expression of SCLC-associated markers (e.g., DLL3), subtype-specific transcription factors (ASCL1, NeuroD1, POU2F3), and/or epithelial markers (e.g., EpCAM, E-cadherin, MUC1). In contrast, clusters 1 and 3 lacked significant expression of these markers and were therefore excluded from the CTC designation (Figure 1D**; Supplementary** Figures 2C).

### 4.10 Differential Abundance Analysis

Observing a potential overrepresentation of cancer cells in multiple clusters, we fit separate mixed-effects logistic regression models for each cluster [45]. In each model, cell condition (normal vs. cancer) was included as a fixed effect, and sample ID was included as a random effect to account for inter-sample variability. From these models, we estimated the odds ratio of a cell in the cluster being cancer versus normal. P values for the condition coefficient were calculated and adjusted for multiple testing (Benjamini-Hochberg), with an adjusted P value < 0.05 considered statistically significant.

### 4.11 Subtype Classification

To classify each cell to the SCLC subtypes within the data, we first subset the data to the three transcription factors reported to distinguish each subtype. We standardized each protein’s expression values using the scale() function with default parameters and performed k-means clustering (k=4) to separate into distinct subtypes. (ComplexHeatmap v2.22.0).

### 4.12 Subtype Treatment Status Association

To identify whether a specific SCLC subtype is associated with a specific treatment status, we fit separate mixed-effects logistic regression models for each subtype. In each model, treatment status (naive vs. CTX+ICI or naive vs. tarlatamab or CTX+ICI vs. tarlatamab) was included as a fixed effect, and patient ID was included as a random effect to account for inter-patient variability. From these models, we estimated the odds of a cell from one treatment group being from a specific subtype relative to the other subtypes. We then reported the odds ratio between the two treatment groups being compared. P values for the treatment status coefficient were calculated and adjusted for multiple testing (Benjamini-Hochberg), with an adjusted P value < 0.05 considered statistically significant.

### 4.13 Down-sampling Subtype Treatment Status Association

To identify whether a. specific SCLC subtype is associated with a specific treatment status, we conducted chi-squared test for each subtype in a subset of equally down-sampled liquid biopsies (30 CTCs per specimen). From these tests, we estimated the odds of a cell from one treatment group being from a specific subtype relative to the other subtypes. We then reported the odds ratio between the two treatment groups being compared. P values were adjusted for multiple testing (Benjamini-Hochberg), with an adjusted P value < 0.05 considered statistically significant.

### 4.14 Violin Plot Statistical Testing

For each violin plot, a two-tailed Wilcoxon rank sum test was used to calculate P values. An adjusted P value of < 0.05 was considered statistically significant.

### 4.15 CTC Percent Calculation

The proportion of CTCs among PBMCs was determined for each patient by dividing the total CTC cluster cell count by the number of non-apoptotic cells in the sample and multiplying the result by 100. The average percentage of CTCs per non-apoptotic events across patients analyzed in our study was 0.0739% (**Supplementary Table 1**).

### 4.16 PHENOSTAMP

#### Reference map and projection model

CyTOF CTC FCS files were projected onto the EMT-MET PHENOtypic STAte MaP (PHENOSTAMP) as previously described by Karacosta *et al* [14]. Conflicts of Interest statement

