## Supplementary File for "CyTOF-based profiling of circulating tumor cells predicts aggressiveness and therapy response in SCLC liquid biopsies at a personalized level"

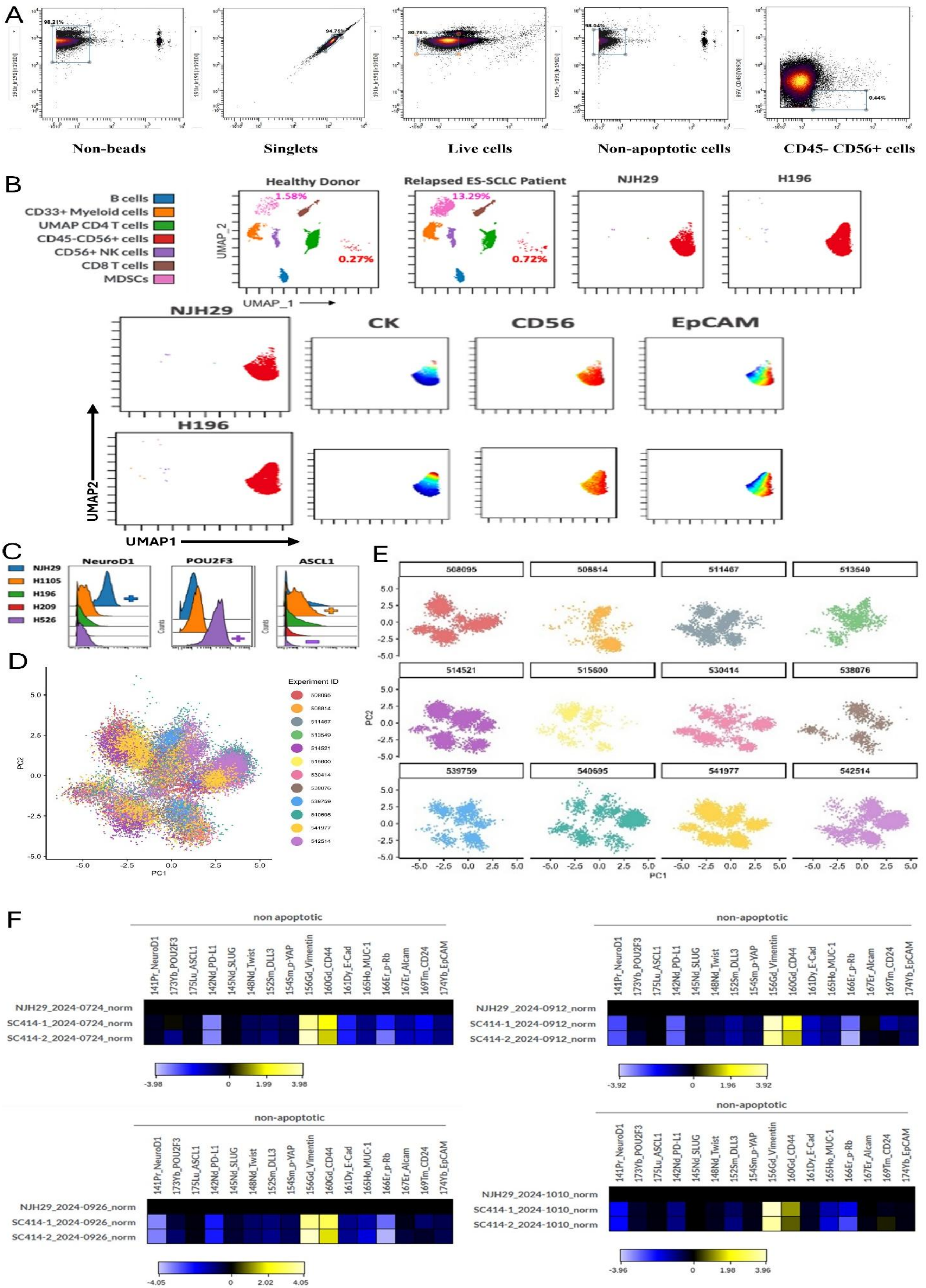
A
98.21%
94.75%
80.78%
98.04%
0.44%
Non-beads
Singlets
Live cells
Non-apoptotic cells
CD45- CD56+ cells
B
B cells
CD33+ Myeloid cells
UMAP CD4 T cells
CD45-CD56+ cells
CD56+ NK cells
CD8 T cells
MDSCs
Healthy Donor
1.58%
0.27%
Relapsed ES-SCLC Patient
13.29%
0.72%
NJH29
H196
UMAP_2
UMAP_1
NJH29
CK
CD56
EpCAM
H196
UMAP2
UMAP1
C
NJH29
H1105
H196
H209
H526
NeuroD1
POU2F3
ASCL1
Counts
D
Experiment ID
508095
508814
511467
513549
514521
515600
530414
538076
539759
540695
541977
542514
PC1
PC2
E
508095
508814
511467
513549
514521
515600
530414
538076
539759
540695
541977
542514
PC1
PC2
F
non apoptotic
non-apoptotic
141Pr_NeuroD1
173Yb_POU2F3
175Lu_ASCL1
142Nd_PD-L1
145Nd_SLUG
148Nd_Twist
152Sm_DLL3
154Sm_p-YAP
156Gd_Vimentin
160Gd_CD44
161Dy_E-Cad
165Ho_MUC-1
166Er_p-Rb
167Er_Alcam
169Tm_CD24
174Yb_EpCAM
NJH29_2024-0724_norm
SC414-1_2024-0724_norm
SC414-2_2024-0724_norm
-3.98
0
1.99
3.98
NJH29_2024-0912_norm
SC414-1_2024-0912_norm
SC414-2_2024-0912_norm
-3.92
0
1.96
3.92
NJH29_2024-0926_norm
SC414-1_2024-0926_norm
SC414-2_2024-0926_norm
-4.05
0
2.02
4.05
NJH29_2024-1010_norm
SC414-1_2024-1010_norm
SC414-2_2024-1010_norm
-3.96
0
1.98
3.96

**Supplementary Figure 1. Detection of CTCs in liquid biopsies using an optimized CyTOF panel with SCLC-specific markers. A.** Gating strategies and hierarchy (left to right): elimination of beads, doublets, dead cells, apoptotic cells and CD45+ cells to obtain CD45-CD56+ populations using Cytobank. **B**. (Top panel) UMAP visualization of PBMCs from healthy donor vs SCLC patient and SCLC cell lines NJH29 and H196; (bottom panel) UMAP visualization of SCLC cells showing expression of pan-Cytokeratin, CD56 and EpCAM. **C.** Histograms showing levels of transcription factors ASCL1, NeuroD1 and POU2F3 in SCLC cell lines H1105 (ASCL1+), NJH29 (NeuroD1+), H526 (POU2F3+), H209 (negative control for POU2F3) and H196 (negative control for ASCL1). **D.** Combined PCA plots of all CyTOF runs from different dates. **E.** PCA plots of all CyTOF samples acquired on different dates demonstrate minimal batch effects, indicating negligible variability across runs. **F.** Heatmaps of arcsinh-transformed marker expression in non-apoptotic cell populations from two samples processed across four different dates, with differential expression normalized to the NJH29 cell line, demonstrating technical replicability of our CyTOF runs.

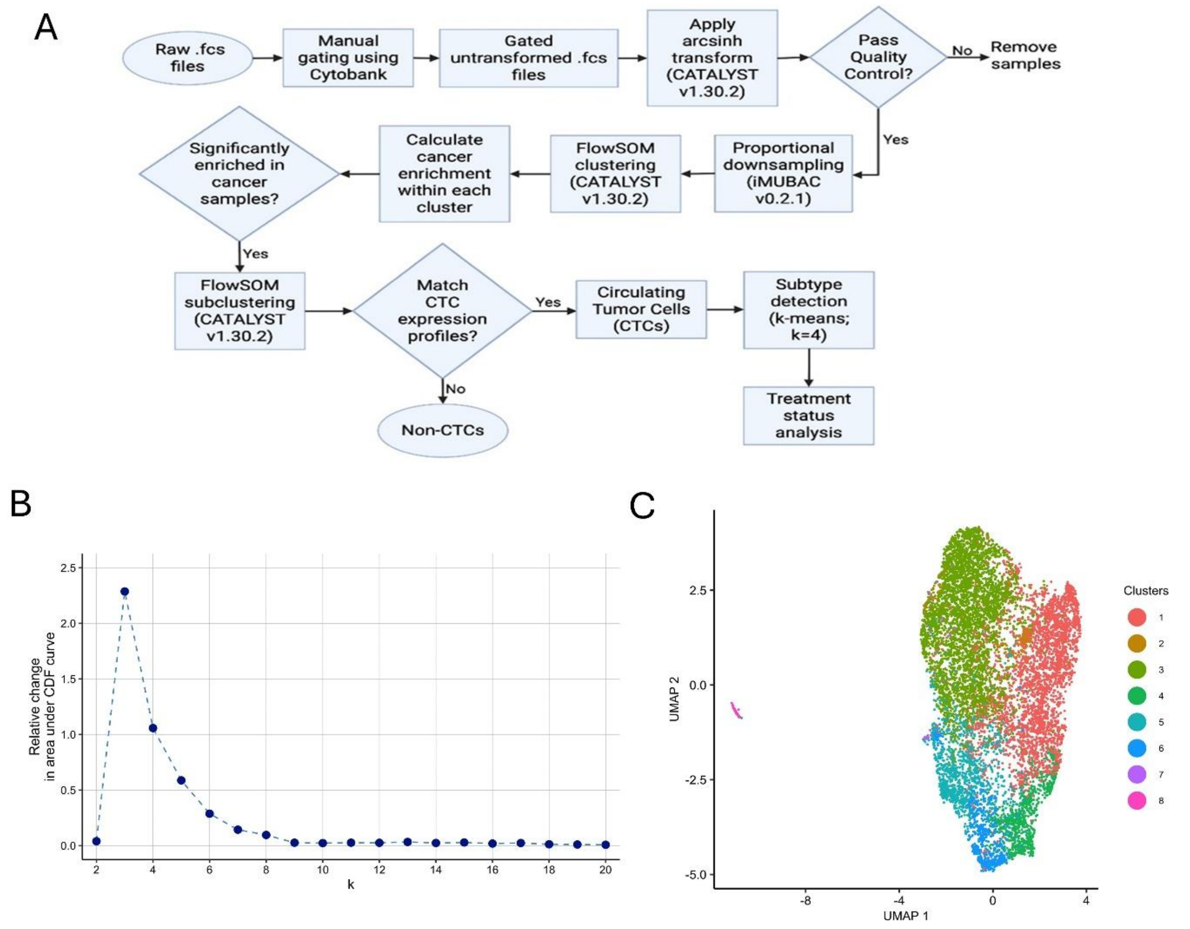


**Supplementary Figure 2. Characterization of CTCs from liquid biopsies using SCLC-specific markers. A.** Flowchart showing the development of the pipeline to detect, characterize and phenotype CTCs from liquid biopsies. **B.** The plot shows the relative change in the area under the cumulative distribution function (CDF) curve (y-axis) as a function of the number of primary clusters (k, x-axis). The plateau in relative change indicates the optimal cluster number (k=8). **C.** UMAP visualization of FlowSOM sub-clustering performed on cancer-enriched primary clusters p6, p7 and p8 from Figure 1C showing eight new secondary clusters (s1-s8).

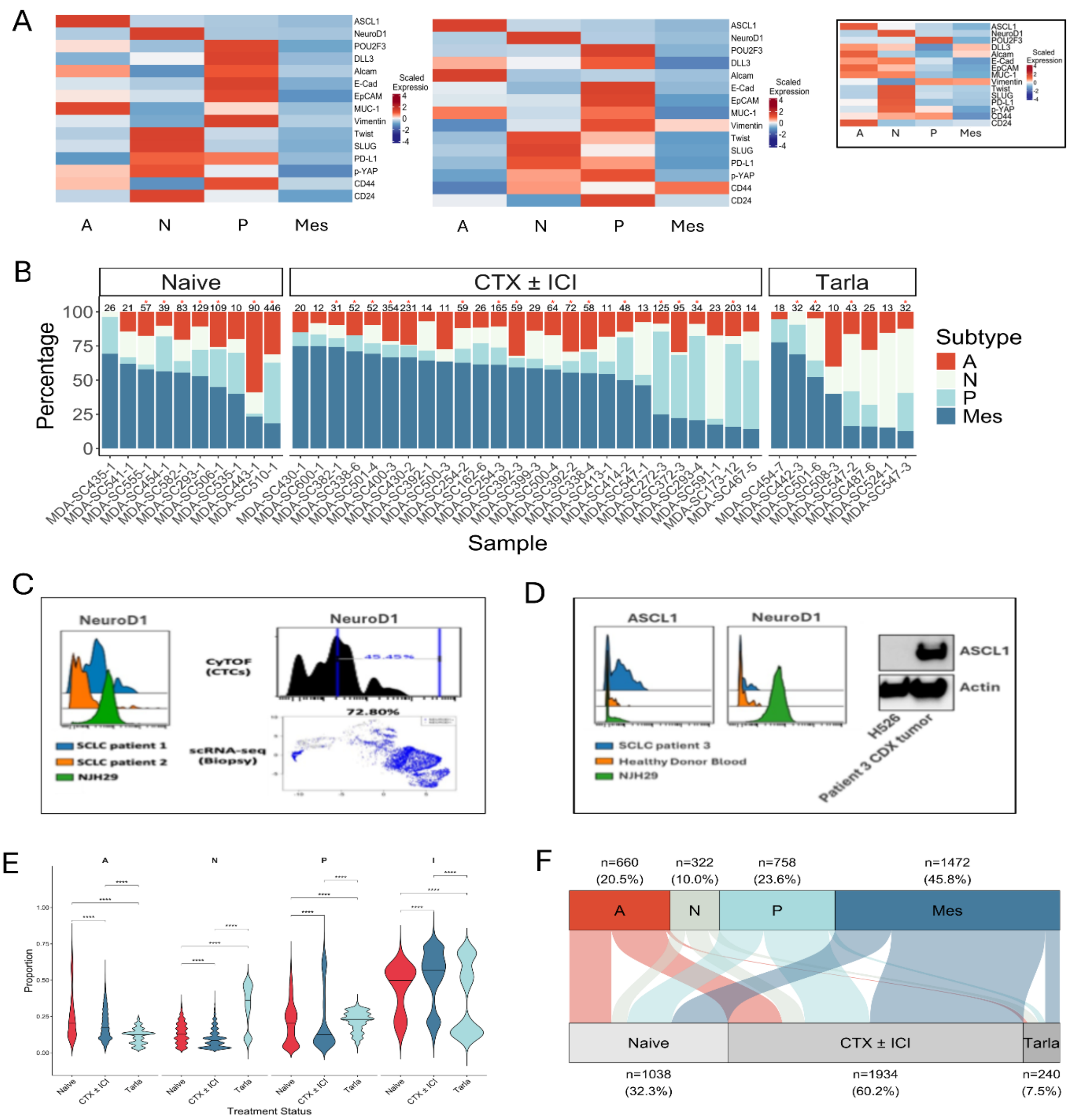


G

| Treatment Status | Subtype | % CTCs |
|---|---|---|
| Naïve | A | 26.5 |
| | N | 9.73 |
| | P | 28.0 |
| | Mes | 35.7 |
| CTX ± ICI | A | 18.5 |
| | N | 7.65 |
| | P | 21.9 |
| | Mes | 52.0 |
| Tarlatamab | A | 11.7 |
| | N | 30.4 |
| | P | 17.9 |
| | Mes | 40 |

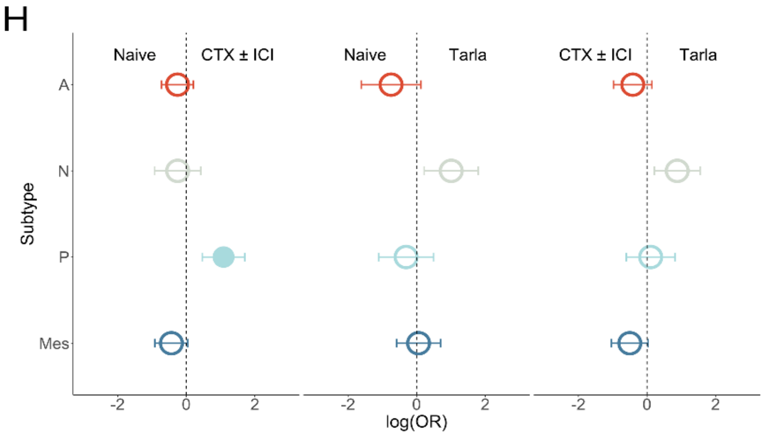

**Supplementary Figure 3. Subtype-specific signatures in CTCs and their treatment associations. A.** (Left) Heatmap showing marker expression profile within each subtype in non-cancer enriched primary clusters p1-p5 from Figure 1B. (Center) Heatmap showing marker expression profile within each subtype in non-CTC secondary clusters s1 and s3 from Supplementary Figure 1C. (Right inset) CTC heatmap from Figure 2 in the main manuscript for comparison. **B.** Bar graphs showing CTC subtype composition across all liquid biopsies in treatment-naïve, CTX ± ICI -treated and tarlatamab treated patients (only liquid biopsies with >=10 CTCs were selected). Red asterisks indicate specimens with at least 30 CTCs detected utilized for the down-sampling analysis related to Figure 2C-E and Supplementary Figure 3E. **C.** (Left) Histograms showing levels of NeuroD1 expression in NJH29 cells (NeuroD1 positive control) and in CD45-CD56+ cells from SCLC patients 1 and 2 as assessed by CyTOF; (right, top) CyTOF NeuroD1 histogram, (right, bottom) single-cell RNA seq NeuroD1 expression levels from matched primary tumor of SCLC patient 1. **D.** (Left) Histograms showing ASCL1 and NeuroD1 expression in NJH29 cells and in CD45-CD56+ cells from a healthy donor and SCLC patient 3 as assessed by CyTOF; (right) western blot showing expression of ASCL1 in CTC-derived xenograft tumor from SCLC patient 3 and negative control H526 cells. **E.** Violin plots showing the distributions of proportions of each subtype across 1000 iterations for random sampling of 30 CTCs per liquid biopsy (related to Figure 2C-E). Comparisons: naive vs CTX ± ICI (P values: A: 6.826e-85, N: 7.84e-293, P: 1.04e-03, Mes: 1.964e-283), naive vs tarlatamab (P values: A: < 0.0001, N: < 0.0001, P: 5.62e-24, Mes: 3.826667e-39), and CTX+ICI vs tarlatamab groups (P values: A: < 0.0001, N: < 0.0001, P: 1.380000e-166, Mes: 4.466667e-280). P values calculated using Wilcoxon rank sum test and adjusted for multiple testing (Benjamini-Hochberg). **F.** Alluvial plot showing the distribution of all CTCs across our entire patient/specimen cohort. Each flow represents the proportion of cells assigned to a subtype within each treatment category. **G.** Table summarizing the percent of CTCs of each subtype in each treatment group in the alluvial plot in F. **H.** Odds ratio (OR) forest plots displaying associations between CTC subtypes and treatment groups (left; naïve vs CTX ± ICI, middle; naive vs tarlatamab, right; CTX ± ICI vs tarlatamab), calculated using mixed-effect logistic regression analysis. Log(OR) > 1 indicates enrichment of a subtype in the right treatment condition relative to all other subtypes, while log(OR) < 1 indicates enrichment of a subtype in the left treatment condition relative to all other subtypes (Naive vs CTX+ICI P values: A: 0.4945, N: 0.4997, P: 0.0029, Mes: 0.2716) (Naive vs tarlatamab P Values: A: 0.2744, N: 0.0486, P: 0.9078, Mes: 0.9078) (CTX ± ICI vs tarlatamab P values: A: 0.3175, N: 0.0380, P: 0.7588, Mes: 0.1815). See Methods for additional details.

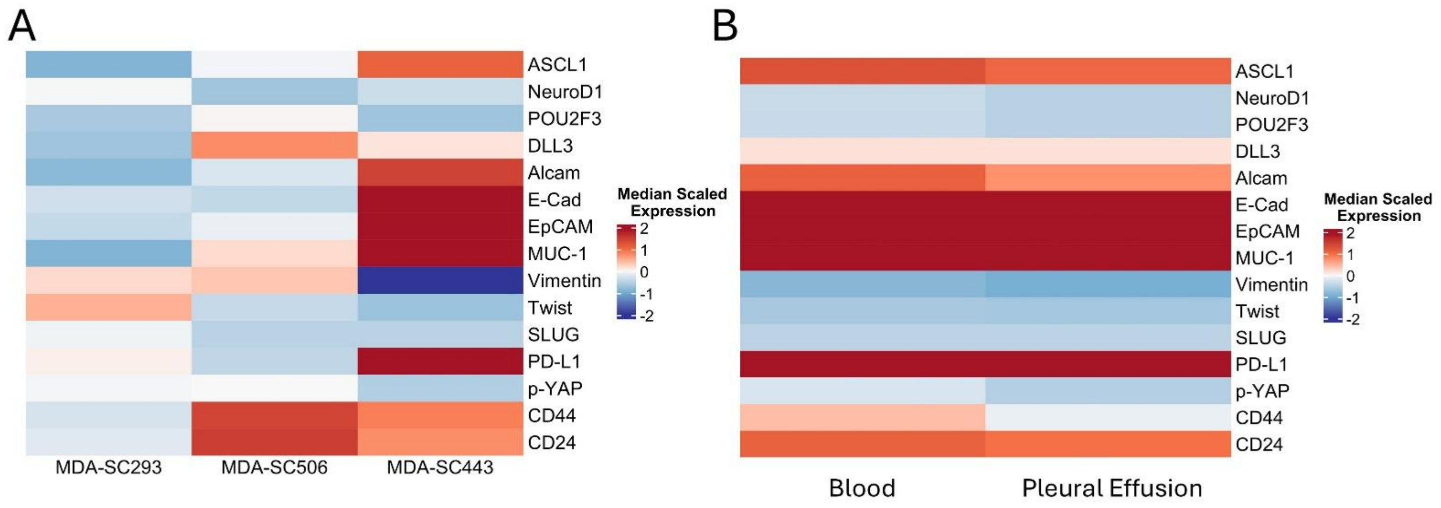


**Supplementary Figure 4. Temporal tracking reveals treatment-associated subtype changes within each patient. A.** Heatmap showing CTC protein expression profiles of the three naïve samples MDA-SC293, MDA-SC506 and MDA-SC443 shown in Figures 3A-D in main text as assessed with CyTOF. **B.** Heatmap showing the near identical expression profile in CD45-CD56+ cells in patient MDA-SC443 blood specimen (left) and pleural effusion (right). ASCL1 high expression was confirmed in both types of liquid biopsies and was in agreement with xenograft tumor IHC shown in Figure 3A.

**Supplementary Table 1. List of Antibodies used for CyTOF**

| Target | Metal label | Clone | Specificities | Source | Catalog number |
|---|---|---|---|---|---|
| CD45 | 89Y | HI30 | Hu, Ch | DVS-Fluidigm | 3089003B |
| CD326, EpCAM | 174Yb | 9C4 | Hu | BioLegend | 324202 |
| CD56 | 139La | NCAM16.2 | Hu | BD | 559043 |
| CD274, PD-L1 | 142Nd | MIH1 | Hu | eBioscience | 14-5983-82 |
| Slug, SNAI2 | 145Nd | S43-1259 | Hu, Ms | BD | 564614 |
| p-Rb | 166Er | J112-906 | Hu, Ms, Rt | DVS-Fluidigm | 3166011A |
| CD24 | 169Tm | ML5 | Hu | DVS-Fluidigm | 3169004B |
| NeuroD1 | 141Pr | R8-294 | Hu, Ms | BD | 563000 |
| p-YAP1 | 154Sm | Polyclonal | Hu | Abcam | ab62751 |
| CD324, E-Caderin | 161Dy | 67A4 | Hu | BioLegend | 324102 |
| Vimentin | 156Gd | RV202 | Hu, Ms, Rb, Rt, cross | DVS-Fluidigm | 3156023A |
| CD44 | 160Gd | IM7 | Hu, Ms, Ch, Rh | BioLegend | 103002 |
| CD166, ALCAM | 167Er | 105902 | Hu | R&D | MAB6561 |
| Twist-1 | 148Nd | Polyclonal | Hu | R&D | AF6230 |
| Caspase 3, cleaved | 153Eu | D3E9 | Hu, Ms, Rt, Bv, Pg | Cell Signaling Technologies | 9579BF |
| TTF-1, NKX2-1 | 164Dy | 2054E | Hu | R&D | MAB94581-100 |
| tumor-MUC1 | 165Ho | N/A | Hu | OncoTAb | HTAB004 |
| Delta-like ligand 3 (DLL3) | 152 Sm | E3J5R | Hu | Cell Signaling Technologies | 88260SF |
| ASCL1 | 175Lu | E5S4Q | Hu | Cell Signaling Technologies | 10585BF |
| POU2F3 | 173Yb | E5N2D | Hu | Cell Signaling Technologies | 38561SF |

*Antibodies to ASCL1, POU2F3, DLL3 and MUC1 were in-house purchased, conjugated and titrated and added at optimal concentrations to cell pellets in addition to the surface and intracellular antibody cocktails provided by the MD Anderson Flow Cytometry Core Facility.

**Supplementary Table 2: CTC percentages in each SCLC patient liquid biopsy**

| sample_id | percent_ctc |
|---|---|
| MDA-SC162-6 | 0.006 |
| MDA-SC173-12 | 0.083 |
| MDA-SC173-12 | 0.007 |
| MDA-SC245-3 | 0.044 |
| MDA-SC254-2 | 0.085 |
| MDA-SC272-3 | 0.162 |
| MDA-SC272-3 | 0.006 |
| MDA-SC293-1 | 0.166 |
| MDA-SC293-4 | 0.631 |
| MDA-SC338-4 | 0.151 |
| MDA-SC338-6 | 0.205 |
| MDA-SC355 | 0 |
| MDA-SC363-5 | 0 |
| MDA-SC370-2 | 0.011 |
| MDA-SC370-2 | 0 |
| MDA-SC372-3 | 0.217 |
| MDA-SC372-3 | 0.003 |
| MDA-SC382-1 | 0.057 |
| MDA-SC392-1 | 0.008 |
| MDA-SC392-1 | 0.002 |
| MDA-SC392-2 | 0.076 |
| MDA-SC392-2 | 0.022 |
| MDA-SC392-3 | 0.061 |
| MDA-SC392-3 | 0.004 |
| MDA-SC399-3 | 0.016 |
| MDA-SC400-1 | 0.107 |
| MDA-SC400-2 | 0 |
| MDA-SC400-3 | 1.001 |
| MDA-SC405-1 | 0.033 |
| MDA-SC405-2 | 0.01 |
| MDA-SC413-1 | 0.036 |
| MDA-SC414-1 | 0.014 |
| MDA-SC414-1 | 0.01 |
| MDA-SC414-2 | 0.063 |
| MDA-SC414-2 | 0.018 |
| MDA-SC414-2 | 0.001 |
| MDA-SC414-9 | 0.031 |
| MDA-SC430-1 | 0.122 |
| MDA-SC430-2 | 0.796 |
| MDA-SC435-1 | 0.054 |
| MDA-SC442-2 | 0.023 |
| MDA-SC442-3 | 0.117 |

| | |
|---|---|
| MDA-SC443-1 | 0.182 |
| MDA-SC454-1 | 0.11 |
| MDA-SC454-1 | 0.003 |
| MDA-SC454-1 | 0.002 |
| MDA-SC454-7 | 0.035 |
| MDA-SC467-5 | 0.019 |
| MDA-SC487-3 | 0.006 |
| MDA-SC487-3 | 0.001 |
| MDA-SC487-4 | 0.009 |
| MDA-SC487-4 | 0.005 |
| MDA-SC487-6 | 0.022 |
| MDA-SC495-4 | 0.001 |
| MDA-SC495-5 | 0.005 |
| MDA-SC499-3 | 0.011 |
| MDA-SC500-3 | 0.019 |
| MDA-SC500-4 | 0.088 |
| MDA-SC500-5 | 0.006 |
| MDA-SC500-5 | 0.002 |
| MDA-SC501-4 | 0.066 |
| MDA-SC501-6 | 0.128 |
| MDAS-C506-1 | 0.226 |
| MDA-SC506-1 | 0.018 |
| MDA-SC508-3 | 0.028 |
| MDA-SC510-1 | 0.635 |
| MDA-SC521-1 | 0.003 |
| MDA-SC524-1 | 0.021 |
| MDA-SC526-2 | 0.001 |
| MDA-SC534-1 | 0.004 |
| MDA-SC535-1 | 0.008 |
| MDA-SC537-1 | 0.002 |
| MDA-SC541-1 | 0.013 |
| MDA-SC541-1 | 0.005 |
| MDA-SC547-1 | 0.103 |
| MDA-SC547-1 | 0.035 |
| MDA-SC547-2 | 0.113 |
| MDA-SC547-2 | 0.015 |
| MDA-SC547-3 | 0.058 |
| MDA-SC547-3 | 0.021 |
| MDA-SC548-1 | 0.006 |
| MDA-SC553-1 | 0.003 |
| MDA-SC555-1 | 0.028 |
| MDA-SC555-1 | 0.014 |
| MDA-SC555-2 | 0.002 |
| MDA-SC571-1 | 0.002 |
| MDA-SC572-1 | 0.003 |
| MDA-SC573-1 | 0.009 |
| MDA-SC574-1 | 0.003 |
| MDA-SC577-1 | 0.003 |

| | |
|---|---|
| MDA-SC582-1 | 0.023 |
| MDA-SC582-1 | 0.014 |
| MDA-SC584-1 | 0.002 |
| MDA-SC591-1 | 0.099 |
| MDA-SC600-1 | 0.03 |

**Supplementary Table 3. Number of patients and CTCs used for each analysis**

| Category | #CTCs for alluvial plot (Figure 2C) from samples that had at least 30 CTCs | #Patients for bar plot (Supplementary Fig 3C) | #Patients for longitudinal tracking (with >=10 CTCs) (Figure 3F-I) |
|---|---|---|---|
| Total | 810 (27 samples from 21 patients) | 44 samples from 33 patients | - |
| Treatment naïve | 210 (7 samples from 7 patients) | 10 samples from 10 patients | - |
| CTX±ICI | 480 (16 samples from 13 patients) | 26 samples from 20 patients | - |
| Tarlatamab | 120 (4 samples from 3 patients) | 8 samples from 7 patients | - |
| Total longitudinally tracked | - | - | 8 |
| Longitudinally tracked (CTX±ICI) | - | - | 5 |
| Longitudinally tracked (tarlatamab) | - | - | 3 |

Note: Some patients overlap between two groups since they had blood drawn at multiple time points and contributed liquid biopsies at both naive and treated stages. Note: One of the 3 tarlatamab-treated patients that were longitudinally tracked was a case of an ALK fusion lung adenocarcinoma transformed to SCLC, and thus not discussed in Figure 3.
